# Mitigating antimicrobial resistance with aspirin (acetylsalicylic acid) and paracetamol (acetaminophen): Conversion of doxycycline and minocycline resistant bacteria into sensitive in presence of aspirin and paracetamol

**DOI:** 10.1101/2021.05.21.445232

**Authors:** Bhoj Raj Singh

## Abstract

The emergence of antimicrobial resistance (AMR) stimulated research for alternatives antimicrobials, repurposing of other drugs, antibiotic adjuvants and alternative therapies for infections. Antimicrobial activity of NSAIDs is often reported and this study evaluated the antimicrobial potential of the two most common NSAIDs, aspirin (acetylsalicylic acid) and paracetamol (acetaminophen), against 293 clinical strains of bacteria. The ability of aspirin and paracetamol to convert minocycline and doxycyclin-resistant bacteria into sensitivity was also tested using micro-broth dilution assays used for determining minimum inhibitory concentration (MIC). Aspirin inhibited all 293 bacterial strains at ≤10.24 mg/ mL concentration. Except for one strain each of Serratia grimaceae and S, aureus paracetamol inhibited none of the 293 strains at 10.24 mg/ mL. Of the 293 strains 116 (39.59%) were sensitive (MIC ≤4 μg/mL) to doxycycline and 127 (43.34%) to minocycline. Of the selected 57 minocycline-resistant (MIC >4 μg/mL) strains aspirin converted 32 (56.14%) to minocycline-sensitive. Of the 49 doxycycline-resistant (MIC >4 μg/mL) strains tested in presence of aspirin 30 (61.22%) turned sensitive. Of the 34 doxycycline-resistant strains tested in presence of paracetamol 11 (32.35%) become sensitive. The study concluded that most of the bacterial strains were not susceptible to aspirin and paracetamol at their concentrations often available in plasma at maximum therapeutic dose levels and had no significant change in their susceptibility to doxycycline and minocycline. The study indicated the potential of aspirin and its combination with antibiotics in the development of therapeutically useful topical antimicrobial formulations.

## Introduction

The global emergence of antimicrobial drug resistance (AMR) even against drugs of the latest classes and generations of antibiotics is a mind-boggling problem for microbiologists and clinicians. Several alternative therapies have been suggested and tried for mitigating the problem of AMR. Scientists have also thought of “Drug repurposing” which is the use of the pre-existing approved drugs for their antimicrobial or antibiotic adjuvant properties (1). The synergism between antibiotics and the repurposed drugs can minimize the therapeutic dose of antibiotics and also the time and costs of the invention of a new drug and putting it for therapeutic use after required approval (2). In the past numerous molecules of non-antibiotic nature earlier acknowledged as anthelmintics, anticancer drugs, antipsychotics, antidepressant drugs, antiplatelets and NSAIDs have been evaluated for their antimicrobial potential (3, 4). Some of the cyclooxygenase inhibitory anti-inflammatory and antipyretic drugs (5) such as acetaminophen (paracetamol), acetylsalicylic acid (aspirin), diclofenac and ibuprofen, flurbiprofen and similar non-steroidal anti-inflammatory drugs (NSAIDs) almost consistently used along with antimicrobial therapy have also been explored for their antimicrobial potential. The most commonly used on-counter NSAIDs (aspirin and paracetamol) are known less for their antibacterial activity in acceptable therapeutic dosages but are reported to enhance the performance of antibiotics either through their synergistic antibacterial action with antibiotics (6–9) or through reducing adherence, production of biofilm, and other virulence factors, and altering antibiotic susceptibility of pathogens (4, 5).

For treatment of *Coronavirus* disease-2019 (COVID-19) two NSAIDs, aspirin (10) and paracetamol (11) along with one or more antibiotics, mainly doxycycline (12) and minocycline (13, 14) are commonly recommended as an adjunct in a COVID-19 treatment regimen. The present study was conducted to evaluate the *in vitro* effect of aspirin on antibacterial activity of doxycycline and minocycline, and of paracetamol on antibacterial activity of doxycycline against selected common potentially pathogenic bacterial strains to understand their interaction.

## Materials and Methods

### Bacterial isolates used in the study

Total 293 bacterial strains of 32 different genera isolated earlier from clinical samples (Tab. 1) and available as glycerol stocks at Clinical Epidemiology Laboratory of Division of Epidemiology, ICAR-Indian Veterinary Research Institutes, Izatnagar were revived and checked for purity and identity (15) were sub-cultured on to nutrient agar (BBL, Difco) slants till tested.

**Table. 1.** The minimum inhibitory concentration (MIC) of aspirin, paracetamol, doxycycline and minocycline for different strains of bacteria.

| Genus of Bacteria | Species of bacteria, numbers of strains tested | Average MIC in µg/ mL (Standard Deviation), resistant strains (MIC≥ 4 µg/ mL |  | Average MIC in mg/ mL (Standard Deviation), resistant strains (MIC≥ 1.28 mg/ mL |  |
| --- | --- | --- | --- | --- | --- |
|  |  | Doxycycline | Minocycline | Aspirin | Paracetamol |
| *Acinetobacter* | *A. alcaligenes* 1, *A. calcoaceticus* 1, *A. lwoffii* 2 | 2.75 (3.5) 1 | 2.38 (3.8) 1 | 1.36 (0.9) 1 | >10.24 |
| *Aerococcus* | *Aerococcus* spp. 2 | 2.5 (2.1) 0 | 0.56 (0.6) 0 | 1.44 (1.6) 1 | >10.24 |
| *Aeromonas* | *A. bestiarum* 5, *A. eucranophila* 2, *A. media* 2, *A. popoffii* 3, *A. salmonicida* 1, *A. schubertii* 1, *A. trota* 2 | 87.45 (141.6) 9 | 81.61 (144.1) 7 | 1.81 (1.2) 7 | >10.24 |
| *Alcaligenes faecalis* | *Alcaligenes faecalis* 1 | 0.50 (NA) 0 | 0.50 (NA) 0 | 0.64 (NA) 0 | >10.24 |
| *Bacillus* | *B. amyloliticus* 1, *B. badius* 1, *B. brevis* 1, *B. cereus* 3, *B. megaterium* 1, *B.* | 9.69 (20.5) 2 | 8.50 (20.9) 1 | 1.72 (0.9) 4 | >10.24 |
|  | *mycoides* 1, *B. sphaericus*<br>1 |  |  |  |  |
| *Brucella* | *B. abortus* 3 | 0.50 (0.0) 0 | 0.50 (0.0) 0 | 0.32 (0.0)<br>0 | >10.24 |
| *Burkholderia* | *B. cepacia* 1 | 512.00<br>(NA) 1 | 128.00<br>(NA) 1 | 10.24<br>(NA) 1 | >10.24 |
| *Edwardsiella* | *E. tarda* 3 | 352.0<br>(277.1) 3 | 182.67<br>(285.6) 2 | 1.07(0.4)<br>0 | >10.24 |
| *Enterobacter* | *E. agglomerans* 10, *E. gregoviae* 1 | 148.64<br>(194.0) 8 | 81.0<br>(148.3) 8 | 2.44 (1.5)<br>6 | >10.24 |
| *Enterococcus* | *E. durans* 1, *E. faecalis* 4,<br>*E. faecium* 5, *E. solitarius* 3 | 250.43<br>(199.2) 12 | 197.86<br>(199.7) 12 | 2.03 (1.5)<br>6 | >10.24 |
| *Erwinia* | *E. amylovora* 2, *E. aphidicola* 1, *E. carotovora* 1, *E. nimipressuralis* 1, *E. stewartii* 1, *E. tasmaniensis* 1 | 76.29<br>(192.1) 2 | 76.29<br>(192.1) 2 | 2.29 (0.7)<br>6 | >10.24 |
| *Escherichia* | *E. coli* 54, *E. fergusonii* 3,<br>*E. hermannii* 1 | 139.19<br>(199.0) 45 | 78.21<br>(144.5) 44 | 2.08 (1.1)<br>29 | >10.24 |
| *Flexibacter* | *Flexibacter* spp. 2 | 1.00 (0.0) 0 | 0.25 (0.0) 0 | 0.64 (0.0)<br>0 | >10.24 |
| *Gallibacterium* | *G. anatis* 2 | 80.00<br>(67.9) 2 | 80.00<br>(67.9) 2 | 2.56 (0.0)<br>2 | >10.24 |
| *Gardnerella* | *Gardnerella* spp. 1 | 512.00<br>(NA) 1 | 32.00 (NA)<br>1 | 1.28 (NA)<br>0 | >10.24 |
| *Geobacillus* | *G. stearothermophilus* 6 | 23.50<br>(51.3) 2 | 21.50<br>(52.2) 1 | 1.92 (1.7)<br>2 | >10.24 |
| *Hafnia* | *H. alvei* 12 | 176.50<br>(215.8) 9 | 124.79<br>(288.1) 9 | 1.97 (0.7)<br>7 | >10.24 |
| *Klebsiella* | *K. oxytoca* 2, *K. pneumoniae* 15 | 162.00<br>(208.9) 16 | 158.38<br>(211.5) 14 | 3.54 (2.2)<br>14 | >10.24 |
| *Kocuria* | *K. rosae* 1 | 0.50 (NA)<br>0 | 0.50 (NA)<br>0 | 0.32 (NA)<br>0 | >10.24 |
| *Micrococcus* | *M. luteus* 1 | 128.00<br>(NA) 1 | 0.25 (NA)<br>0 | 0.32 (NA)<br>0 | >10.24 |
| *Moellerella* | *M. wisconsensis* 1 | 128.00<br>(NA) 1 | 1.00 (NA)<br>0 | 1.28 (NA)<br>0 | >10.24 |
| *Moraxella* | *M. bovis* 1, *M. osloensis* 1,<br>*M. phenylpyruvica* 1 | 6.33 (8.4) 1 | 6.08 (8.6) 1 | 1.81 (1.3)<br>2 | >10.24 |
| *Paenibacillus* | *P. lactis* 1, *P. larvae* 1, *P. pantothenicus* 13 | 13.41<br>(32.3) 6 | 2.27 (4.3) 2 | 1.00 (0.8)<br>3 | >10.24 |
| *Pasteurella* | *P. canis* 2, *P. multocida* 1 | 2.71 (2.2) 0 | 2.71 (2.2) 0 | 1.07 (1.3)<br>1 | >10.24 |
| *Proteus* | *P. mirabilis* 5, *P. penneri*<br>1, *P. vulgaris* 1 | 338.29.0<br>(217.7) 7 | 110.29<br>(182.1) 6 | 2.19 (1.4)<br>3 | >10.24 |
| *Pseudomonas* | *P. aeruginosa* 5, *P.* | 220.57 | 165.71 | 5.85 (4.3) | >10.24 |
|  | *paucimobilis* 1, *P. pseudoalcaligenes* 1 | (203.8) 7 | (159.1) 7 | 5 |  |
| *Raoultella* | *R. terrigena* 6 | 90.17<br>(128.6) 4 | 52.83<br>(100.1) 4 | 2.88 (1.9)<br>4 | >10.24 |
| *Salmonella* | *S. kentucky* 1, *S. Typhimurium* 3, *S. Virchow* 1 | 124.80<br>(217.7) 5 | 24.00<br>(24.7) 5 | 2.56 (1.6)<br>3 | >10.24 |
| *Serratia* | *S. fonticola* 1, *S. grimeae* 2, *S. marcescens* 2, *S. odorifera* 2 | 149.29<br>(247.8) 3 | 149.29<br>(247.8) 3 | 2.56 (0.0)<br>7 | >10.24* |
| *Staphylococcus* | *S. arlettae* 1, *S. aureus* 9, *S. capitis* ssp. *capitis* 3, ssp. *urealyticus* 1, *S. caseolytus* 1, *S. chromogenes* 2, *S. delphini* 4, *S. epidermidis* 10, *S. equorum* 1, *S. gallinarum* 1, *S. haemolyticus* 6, *S. hominis* 6, *S. hyicus* 1, *S. lentus* 1, *S. lugdunensis* 4, *S. schleiferi* 1, *S. sciuri* 1, *S. xylosus* 1 | 48.67<br>(108.3) 21 | 135.57<br>(208.5) 26 | 3.72 (3.6)<br>29 | >10.24* |
| *Streptococcus* | *S. milleri* 5, *S. pneumoniae* | 169.19 | 88.2 | 2.17 (1.5) | >10.24 |
|  | 2, *S. porcinus* 1, *S.*<br>*pyogenes* 1, *S. suis* 3, *S.*<br>*salivaris* 1 | (238.9) 7 | (188.9) 6 | 6 |  |
| *Xenorhabdus* | *X. bovienni* 1 | 8.00 (NA)<br>1 | 8.00 (NA)<br>1 | 2.56 (NA)<br>1 | >10.24 |
| Total | 293 | 115.87<br>(182.46)<br>177 | 93.05<br>(172.20)<br>166 | 2.47<br>(2.27) 150 | NA<br>(NA)<br>293 |

### Determination of minimum inhibitory concentration (MIC) of NSAIDs and antibiotics

The MIC was determined using the micro-broth dilution method in 96 well plates following the CLSI (16, 17) guidelines.

*Stock Solutions:*

1. Aspirin (acetylsalicylic acid, Sigma Aldrich, USA) was dissolved in ethanol at a concentration of 40.96 mg/ mL (4×) and stored at 4-8°C as stock solution till tested.
2. Paracetamol (acetaminophen, Sigma) was dissolved in dimethylsulfoxide (DMSO, Sigma) at a concentration of 20.48 mg/ mL (2×) and stored at 4-8°C as stock solution till tested.
3. Doxycycline hydrochloride hemiethanolate hemihydrates (Sigma) was dissolve in sterile distilled water to have 10.24 mg doxycyline / mL of solution (20×), filter sterilized using 0.2 micron syringe filter and stored at 4-8°C as stock solution till tested.
4. Minocycline (Sigma) was dissolved in sterile distilled water to have 20.48 mg doxycyline / mL (20×), filter sterilized using 0.2 micron syringe filter and stored at 4-8°C as stock solution till tested.

Throughout the study for broth culture of bacteria and for determining MIC Mueller Hinton broth (MHB) medium (BBL Difco) was used. For determining MIC, the test strain of bacteria was grown in MHB at 37°C to the required period (6-8 h) to obtain culture density of about 0.5 OD_590_. Cultures were kept on ice till used for MIC testing within 24 h. For determining MIC two-fold serial dilutions of the test compounds (aspirin and paracetamol starting from 10.24 mg/mL; doxycycline starting from 512 μg/ mL, minocycline starting from 1024 μg/ mL) were made in MHB in 150μL in each of the 96 wells of sterile culture plates (Tarson India Ltd.). In 96 well plates having suitably diluted compound to be tested 1.5 μL of test culture was dispensed aseptically in three rows of the test plate. Then 2^nd^ culture was dispended in the next three rows keeping one empty row as control (to check any contamination in testing). In each plate, for each culture, one column was kept without any antimicrobial compound as positive (for bacterial growth) control. The lid was applied on the plates and plates were incubated at 37°C for 24 h and then growth (opacity) was read using a plate reader at 590nm. The last dilution with no readable growth was recorded as the MIC of the test compound.

### Determination of the effect of aspirin/ paracetamol on antimicrobial activity of doxycycline/ minocycline

The MIC of doxycycline/ minocycline in presence of aspirin/ paracetamol was determined for selected bacterial strains using the similar microdilution method described above for individual drug MIC. The difference was that instead of using plain MHB for making antibiotic’s dilution aspirin/ paracetamol dilutions were made vertically (columns) starting from 1.28 mg/mL to 0.01 mg/mL and then serial two-fold antibiotic’s dilutions were made in rows (horizontally) starting from 512 μg/ mL. Plates were culture inoculated as above for MIC and growth of bacteria was determined to record the no-growth wells for each row. The effect of aspirin was determined both on the MIC of minocycline (for 165 strains, Tab. 2) and doxycycline (for 149 strains, Tab. 3) while the effect of paracetamol on MIC was determined on doxycycline only for 113 strains of bacteria (Tab. 4).

**Table. 2.** The minimum inhibitory concentration (MIC) minocycline for different strains of bacteria in presence of aspirin.

| Genus of bacteria tested | Species of bacteria tested and their numbers | Minimum concentration of aspirin (mg/ mL) having MIC decreasing effect | Percent decrease in MIC of minocycline in presence of aspirin (mg/ mL) |  |  |  |  |  |  |  |
| --- | --- | --- | --- | --- | --- | --- | --- | --- | --- | --- |
|  |  |  | 1.28 | 0.64 | 0.32 | 0.16 | 0.08 | 0.04 | 0.02 | 0.01 |
| *Acinetobacter* | *A. alcaligenes* 1, *A. lwoffii* 2 | 0.04 | 0.94 | 0.93 | 0.93 | 0.91 | 0.64 | 0.34 | 0.00 | 0.00 |
| *Aeromonas* | *A. bestiarum* 1, *A. eucranophila* 2, *A. media* 2, *A. popoffi* 2, *A. scubertii* 2 | 1.28 | 0.18 | 0.00 | 0.00 | 0.00 | 0.00 | 0.00 | 0.00 | 0.00 |
| *Alcaligenes* | *Alcaligenes faecalis* | 0.64 | 0.75 | 0.50 | 0.00 | 0.00 | 0.00 | 0.00 | 0.00 | 0.00 |
| *Bacillus* | *B. brevis* 1, *B. cereus* 2, | NA | -24.70 | -18.30 | -24.70 | -24.70 | -24.70 | -18.30 | -15.10 | -13.50 |
|  | *B. megaterium* 1, *B. mycoides* 1 |  |  |  |  |  |  |  |  |  |
| *Burkholderia* | *Burkholderia cepacia* 1 | NA | 0.00 | 0.00 | 0.00 | 0.00 | 0.00 | 0.00 | 0.00 | 0.00 |
| *Edwardsiella tarda* | *Edwardsiella tarda* 1 | 0.02 | 0.99 | 0.98 | 0.97 | 0.93 | 0.86 | 0.78 | 0.78 | 0.00 |
| *Enterobacter* | *E. agglomerans* 5, *E. gregoviae* 1 | 0.01 | 0.97 | 0.94 | 0.88 | 0.98 | 0.97 | 0.97 | 0.95 | 0.93 |
| *Enterococcus* | *E. faecalis* 3, *E. faecium* 6, *E. malodoratus* 1, *E. solitarius* 2 | 0.02 | 0.99 | 0.99 | 0.99 | 0.98 | 0.96 | 0.87 | 0.83 | 0.00 |
| *Escherichia* | *E. coli* 25, *E. fergusonii* 3, *E. hermani* 1 | 0.01 | 0.91 | 0.85 | 0.71 | 0.74 | 0.72 | 0.69 | 0.66 | 0.55 |
| *Flexibacter* | *Flexibacter* spp. 2 | 0.04 | 0.89 | 0.78 | 0.69 | 0.56 | 0.50 | 0.25 | 0.00 | 0.00 |
| *Geobacillus* | *Geobacillus stearothermophilus* 3 | NA | 0.00 | 0.00 | 0.00 | 0.00 | 0.00 | 0.00 | 0.00 | 0.00 |
| *Hafnia* | *Hafnia alvei* 6 | 0.01 | 0.32 | 0.31 | 0.29 | 0.20 | 0.20 | 0.20 | 0.20 | 0.20 |
| *Klebsiella* | *Klebsiella pneumoniae*<br>ssp. *pneumoniae* 12 | 0.16 | 0.68 | 0.29 | 0.18 | 0.11 | 0.00 | 0.00 | 0.00 | 0.00 |
| *Kocuria rosea* | *Kocuria rosea* | 0.64 | 0.75 | 0.50 | 0.00 | 0.00 | 0.00 | 0.00 | 0.00 | 0.00 |
| *Micrococcus* | *Micrococcus luteus* 1 | 1.28 | 0.50 | 0.00 | 0.00 | 0.00 | 0.00 | 0.00 | 0.00 | 0.00 |
| *Moelerella* | *Moelerella wisconsensis*<br>1 | NA | 0.00 | 0.00 | 0.00 | 0.00 | 0.00 | 0.00 | 0.00 | 0.00 |
| *Moraxella* | *M. bovis* 1, *M. ovis* 1 | 0.64 | 0.50 | 0.50 | 0.00 | 0.00 | 0.00 | 0.00 | 0.00 | 0.00 |
| *Paenibacillus* | *P. lactis* 1, *P. larvae* 1, *P.*<br>*pantothenicus* 10 | 0.64 | 0.50 | 0.50 | 0.00 | 0.00 | 0.00 | 0.00 | 0.00 | 0.00 |
| *Proteus* | *P. mirabilis* 5, *P. penneri*<br>1, *P. vulgaris* 1 | 0.64 | 0.20 | 0.13 | 0.00 | 0.00 | 0.00 | 0.00 | 0.00 | 0.00 |
| *Pseudomonas* | *P. aeruginosa* 5, *P.*<br>*paucimobilis* 1, *P.*<br>*pseudoalcaligenes* 1 | 0.16 | 0.20 | 0.19 | 0.17 | 0.04 | 0.00 | 0.00 | 0.00 | 0.00 |
| *Raoultella* | *Raoultella terrigena* 3 | 0.01 | 0.54 | 0.25 | 0.21 | 0.14 | 0.14 | 0.14 | 0.14 | 0.14 |
| *Salmonella* | *S. Kentucky* 1, *S.* | 0.64 | 0.52 | 0.20 | 0.00 | 0.00 | 0.00 | 0.00 | 0.00 | 0.00 |
|  | *Naestwed* 1, *S.*<br>*Typhimurium* 2 |  |  |  |  |  |  |  |  |  |
| *Staphylococcus* | *S. aureus* 4, *S. capitis*<br>ssp. *capitis* 1, ssp.<br>*urealyticus* 1, *S.*<br>*caseolytus* 1, *S.*<br>*chromogenes* 1, *S.*<br>*delphini* 4, *S. epidermidis*<br>3, *S. gallinarum* 1, *S.*<br>*haemolyticus* 4, *S.*<br>*hominis* 2, *S. hyicus* 1, *S.*<br>*lentus* 1, *S. lugdunensis* 1,<br>*S. sciuri* 1, *S. xylosus* 1 | 0.01 | 0.96 | 0.90 | 0.77 | 0.87 | 0.87 | 0.86 | 0.86 | 0.86 |
| *Streptococcus* | *S. milleri* 3, *S.*<br>*pneumoniae* 2, *S.*<br>*pyogenes* 1, *S. salivaris* 1 | 0.02 | 0.84 | 0.83 | 0.75 | 0.67 | 0.67 | 0.42 | 0.08 | 0.00 |

**Table. 3.**
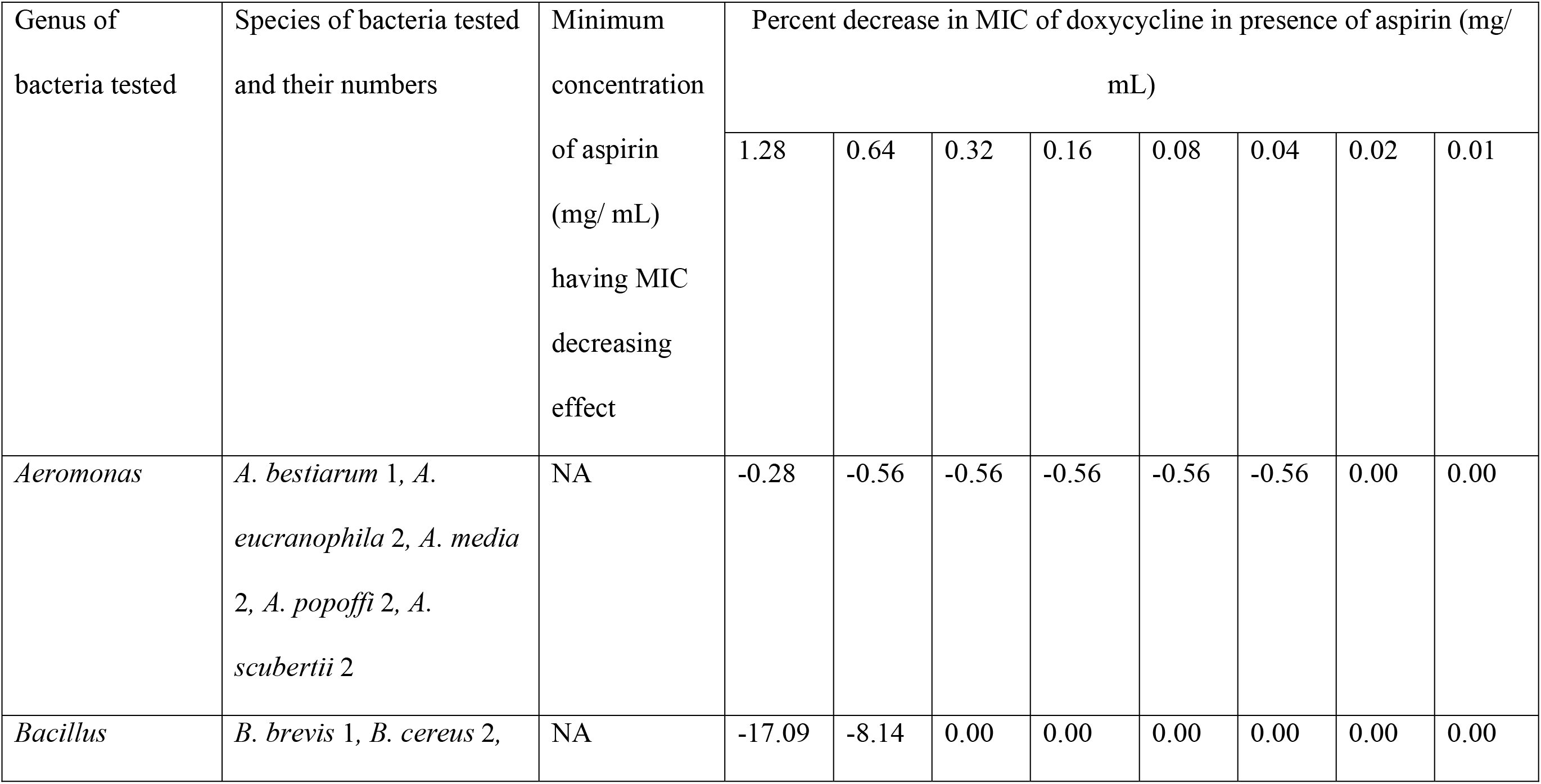
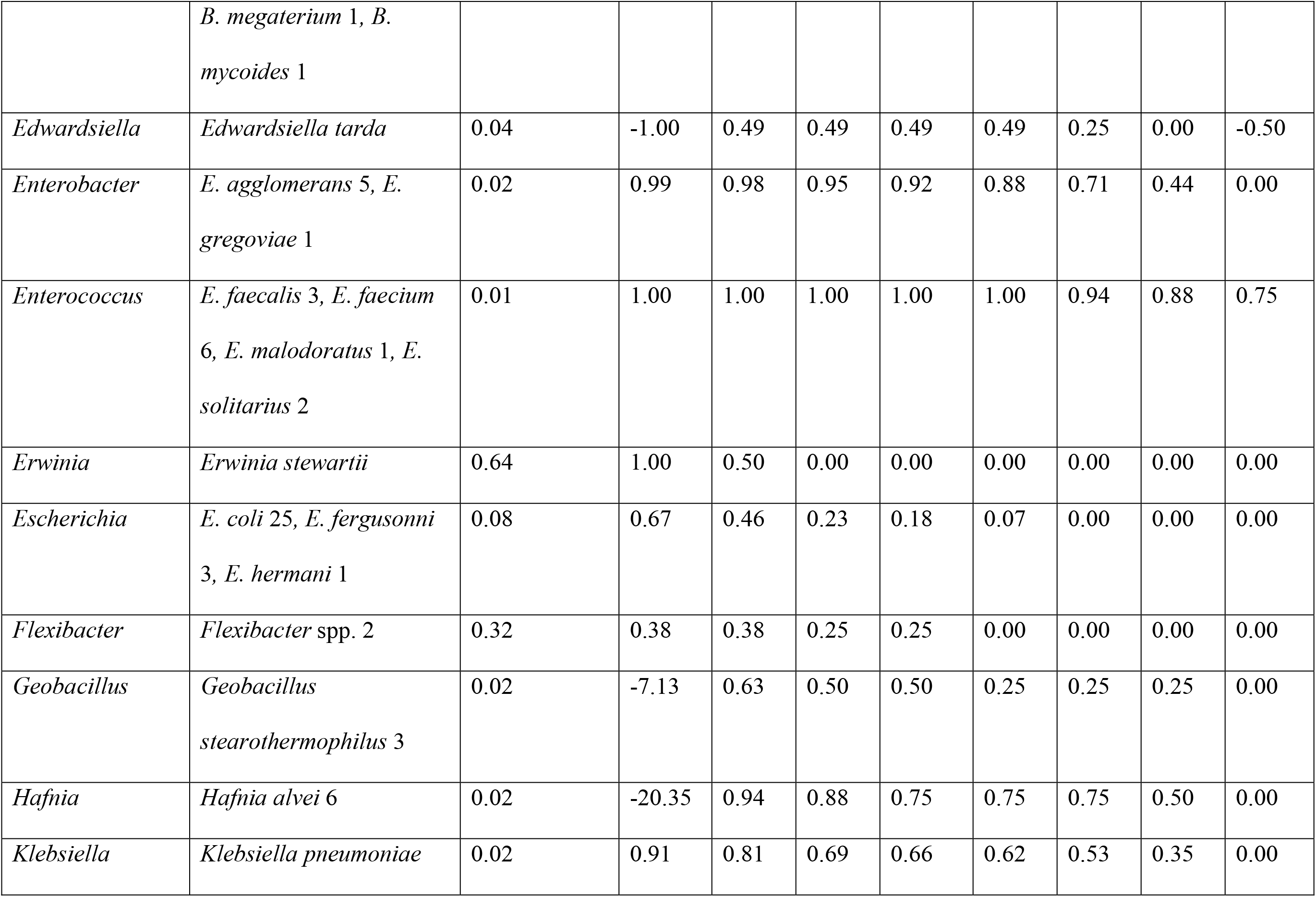
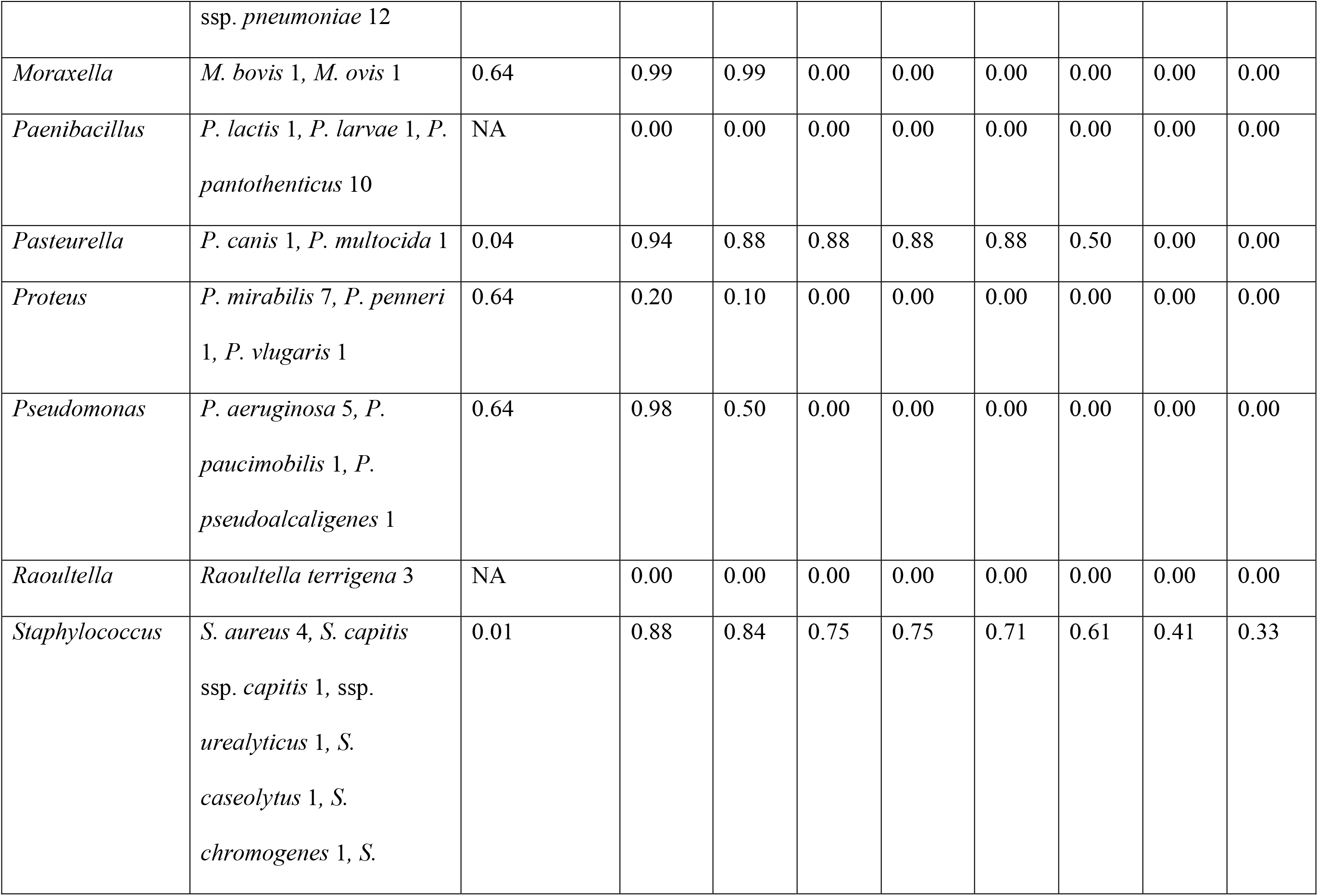
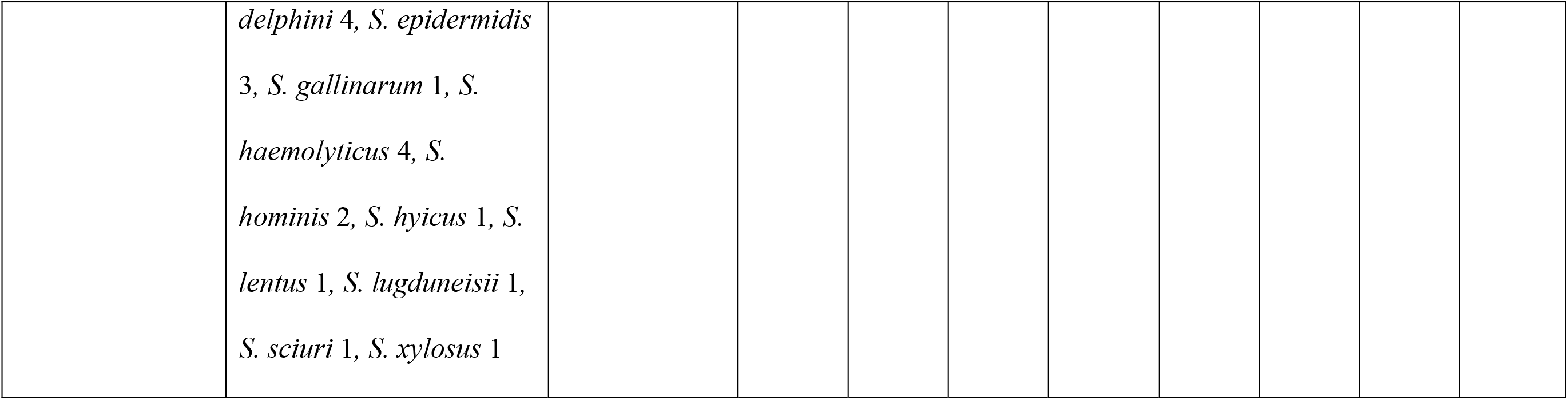
The minimum inhibitory concentration (MIC) doxycycline for different strains of bacteria in presence of aspirin.

| Genus of<br>bacteria tested | Species of bacteria tested<br>and their numbers | Minimum<br>concentration<br>of aspirin<br>(mg/ mL)<br>having MIC<br>decreasing<br>effect | Percent decrease in MIC of doxycycline in presence of aspirin (mg/<br>mL) |  |  |  |  |  |  |  |
| --- | --- | --- | --- | --- | --- | --- | --- | --- | --- | --- |
|  |  |  | 1.28 | 0.64 | 0.32 | 0.16 | 0.08 | 0.04 | 0.02 | 0.01 |
| *Aeromonas* | *A. bestiarum* 1, *A. eucranophila* 2, *A. media* 2, *A. popoffi* 2, *A. scubertii* 2 | NA | -0.28 | -0.56 | -0.56 | -0.56 | -0.56 | -0.56 | 0.00 | 0.00 |
| *Bacillus* | *B. brevis* 1, *B. cereus* 2, | NA | -17.09 | -8.14 | 0.00 | 0.00 | 0.00 | 0.00 | 0.00 | 0.00 |
|  | *B. megaterium* 1, *B. mycoides* 1 |  |  |  |  |  |  |  |  |  |
| *Edwardsiella* | *Edwardsiella tarda* | 0.04 | -1.00 | 0.49 | 0.49 | 0.49 | 0.49 | 0.25 | 0.00 | -0.50 |
| *Enterobacter* | *E. agglomerans* 5, *E. gregoviae* 1 | 0.02 | 0.99 | 0.98 | 0.95 | 0.92 | 0.88 | 0.71 | 0.44 | 0.00 |
| *Enterococcus* | *E. faecalis* 3, *E. faecium* 6, *E. malodoratus* 1, *E. solitarius* 2 | 0.01 | 1.00 | 1.00 | 1.00 | 1.00 | 1.00 | 0.94 | 0.88 | 0.75 |
| *Erwinia* | *Erwinia stewartii* | 0.64 | 1.00 | 0.50 | 0.00 | 0.00 | 0.00 | 0.00 | 0.00 | 0.00 |
| *Escherichia* | *E. coli* 25, *E. fergusonii* 3, *E. hermani* 1 | 0.08 | 0.67 | 0.46 | 0.23 | 0.18 | 0.07 | 0.00 | 0.00 | 0.00 |
| *Flexibacter* | *Flexibacter* spp. 2 | 0.32 | 0.38 | 0.38 | 0.25 | 0.25 | 0.00 | 0.00 | 0.00 | 0.00 |
| *Geobacillus* | *Geobacillus stearothermophilus* 3 | 0.02 | -7.13 | 0.63 | 0.50 | 0.50 | 0.25 | 0.25 | 0.25 | 0.00 |
| *Hafnia* | *Hafnia alvei* 6 | 0.02 | -20.35 | 0.94 | 0.88 | 0.75 | 0.75 | 0.75 | 0.50 | 0.00 |
| *Klebsiella* | *Klebsiella pneumoniae* *ssp. pneumoniae* 12 | 0.02 | 0.91 | 0.81 | 0.69 | 0.66 | 0.62 | 0.53 | 0.35 | 0.00 |
| *Moraxella* | *M. bovis* 1, *M. ovis* 1 | 0.64 | 0.99 | 0.99 | 0.00 | 0.00 | 0.00 | 0.00 | 0.00 | 0.00 |
| *Paenibacillus* | *P. lactis* 1, *P. larvae* 1, *P. pantothenicus* 10 | NA | 0.00 | 0.00 | 0.00 | 0.00 | 0.00 | 0.00 | 0.00 | 0.00 |
| *Pasteurella* | *P. canis* 1, *P. multocida* 1 | 0.04 | 0.94 | 0.88 | 0.88 | 0.88 | 0.88 | 0.50 | 0.00 | 0.00 |
| *Proteus* | *P. mirabilis* 7, *P. penneri* 1, *P. vulgaris* 1 | 0.64 | 0.20 | 0.10 | 0.00 | 0.00 | 0.00 | 0.00 | 0.00 | 0.00 |
| *Pseudomonas* | *P. aeruginosa* 5, *P. paucimobilis* 1, *P. pseudoalcaligenes* 1 | 0.64 | 0.98 | 0.50 | 0.00 | 0.00 | 0.00 | 0.00 | 0.00 | 0.00 |
| *Raoultella* | *Raoultella terrigena* 3 | NA | 0.00 | 0.00 | 0.00 | 0.00 | 0.00 | 0.00 | 0.00 | 0.00 |
| *Staphylococcus* | *S. aureus* 4, *S. capitis* ssp. *capitis* 1, ssp. *urealyticus* 1, *S. caseolytus* 1, *S. chromogenes* 1, *S.* | 0.01 | 0.88 | 0.84 | 0.75 | 0.75 | 0.71 | 0.61 | 0.41 | 0.33 |
|  | *delphini* 4, *S. epidermidis*<br>3, *S. gallinarum* 1, *S.*<br>*haemolyticus* 4, *S.*<br>*hominis* 2, *S. hyicus* 1, *S.*<br>*lentus* 1, *S. lugdunensis* 1,<br>*S. sciuri* 1, *S. xylosus* 1 |  |  |  |  |  |  |  |  |  |

**Table. 4.**
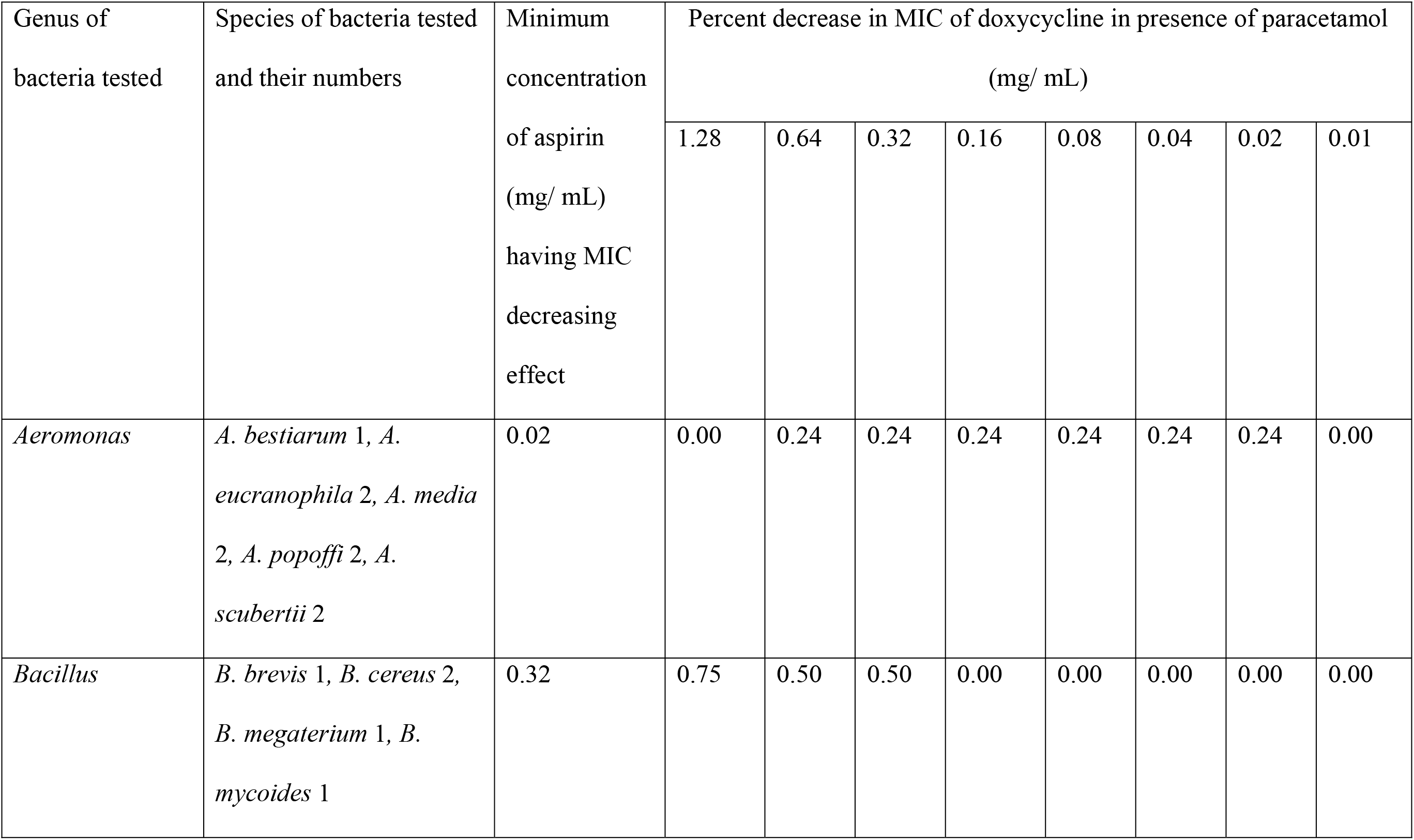
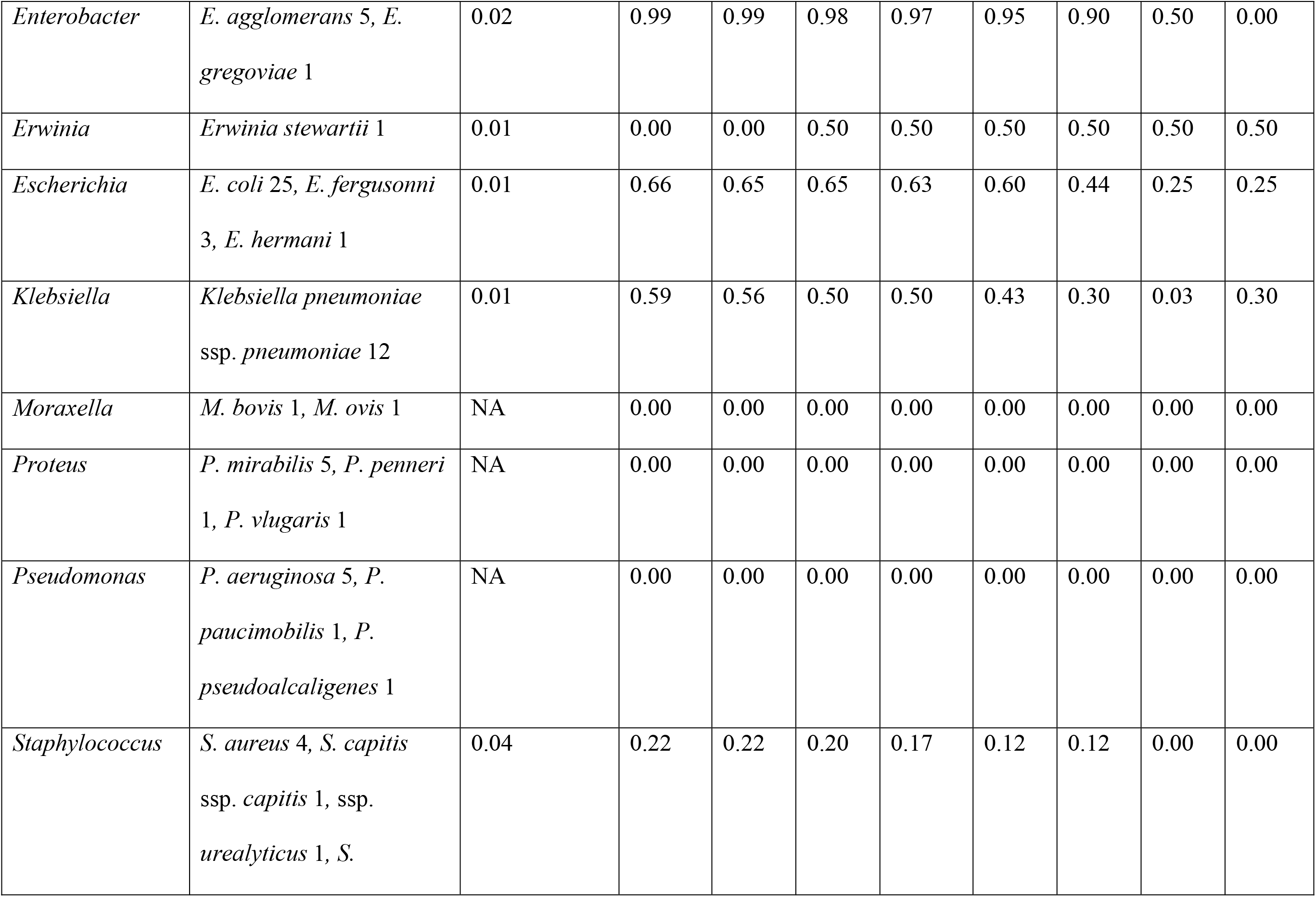
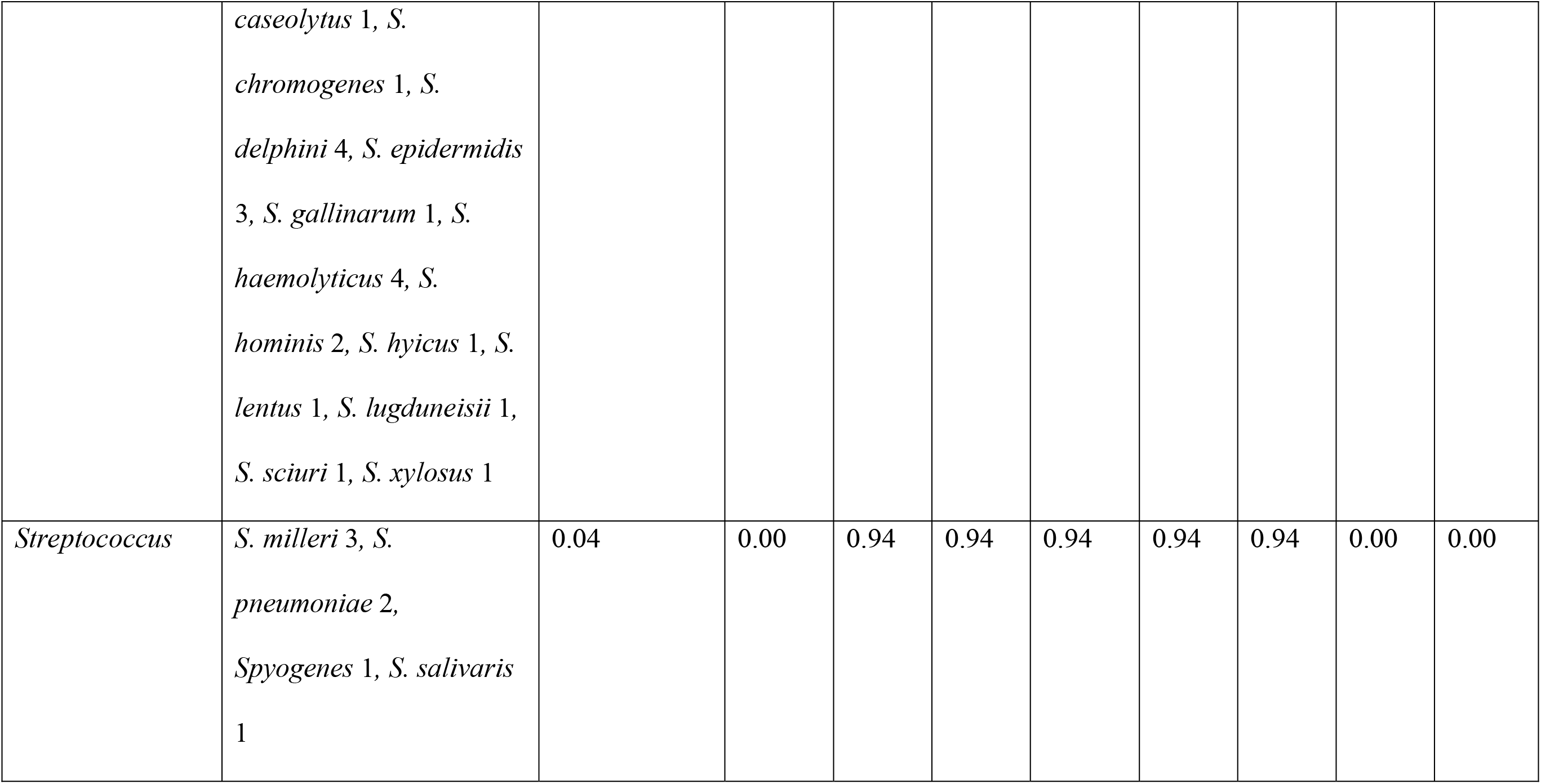
The minimum inhibitory concentration (MIC) doxycycline for different strains of bacteria in presence of paracetamol.

**Table 5.**
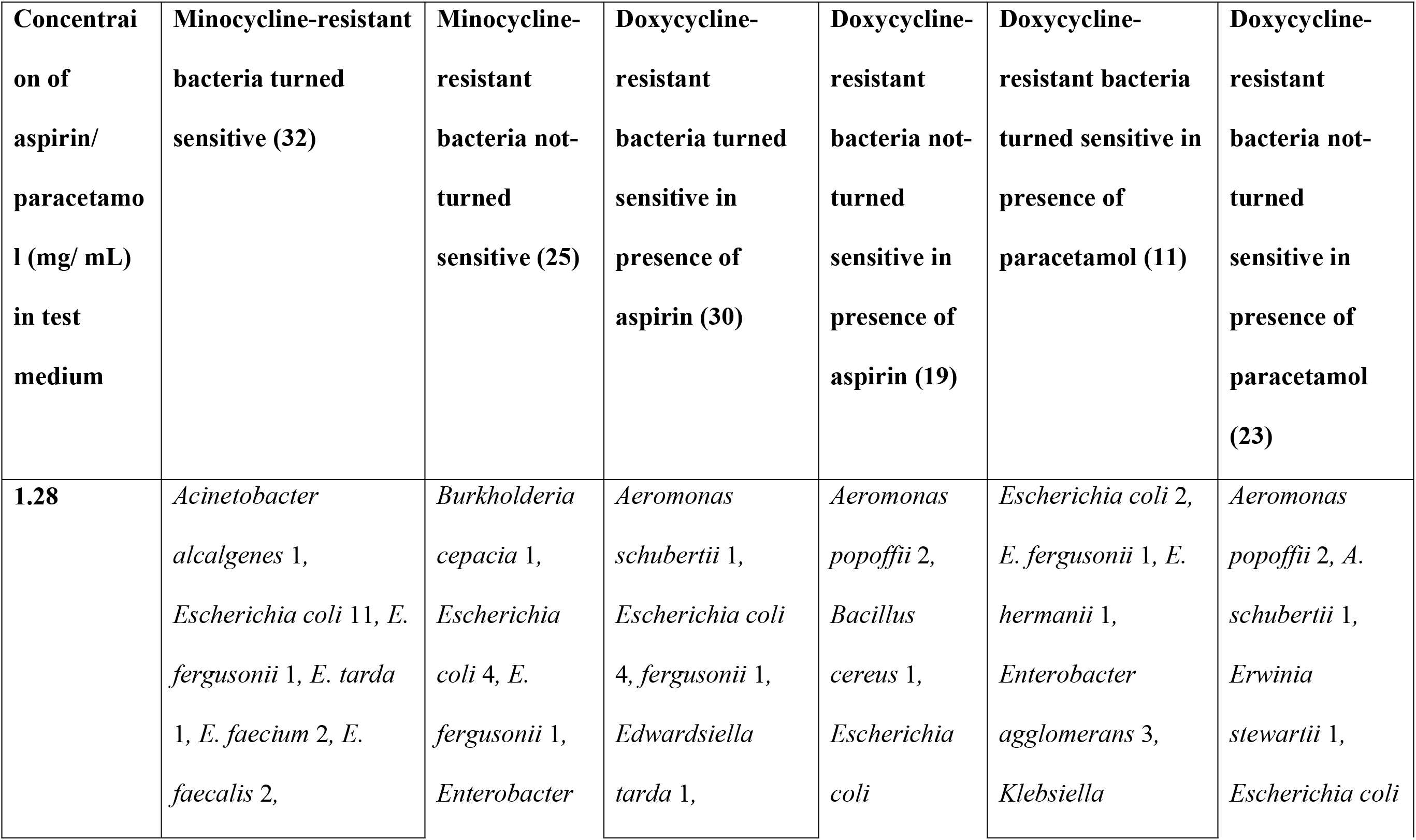
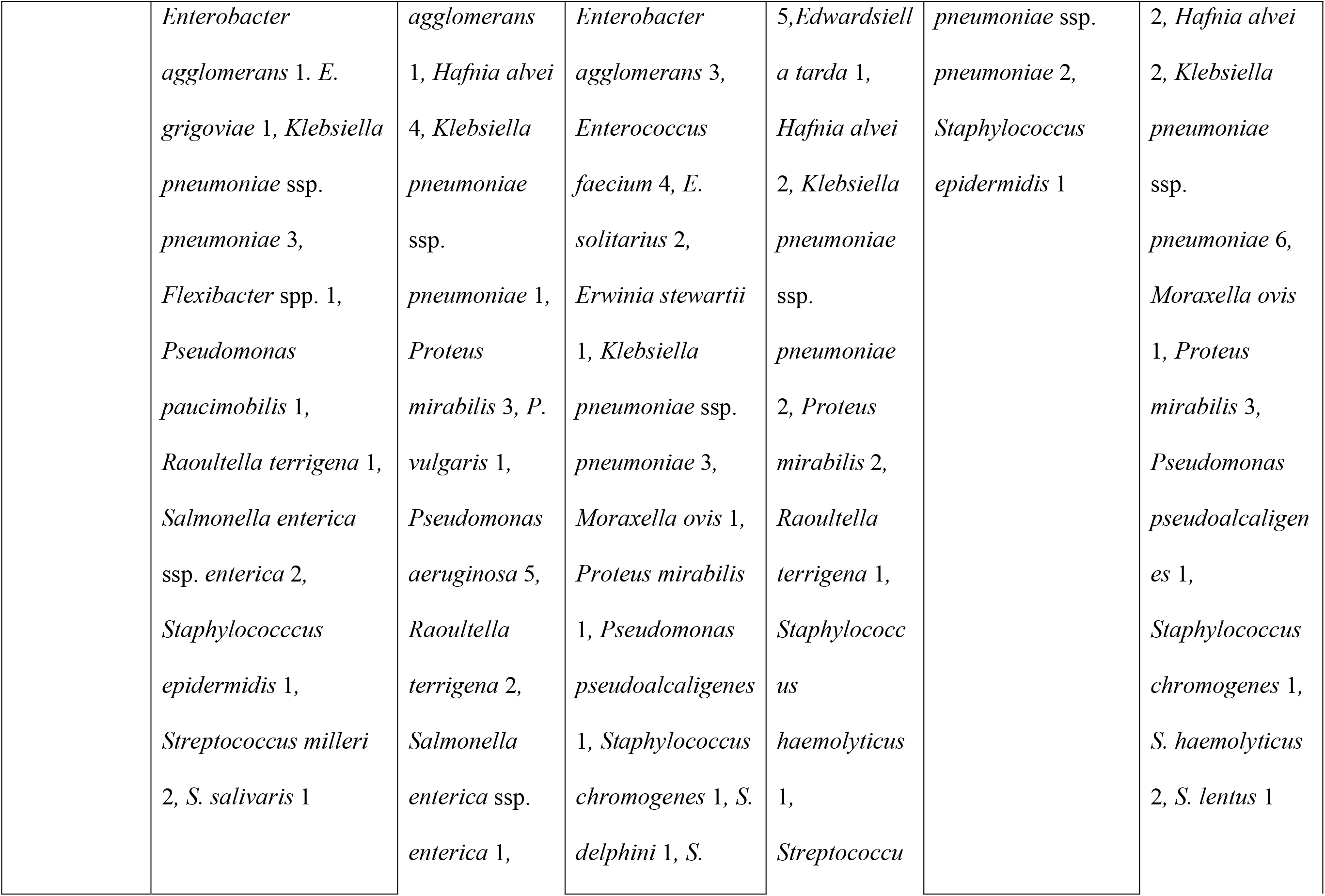
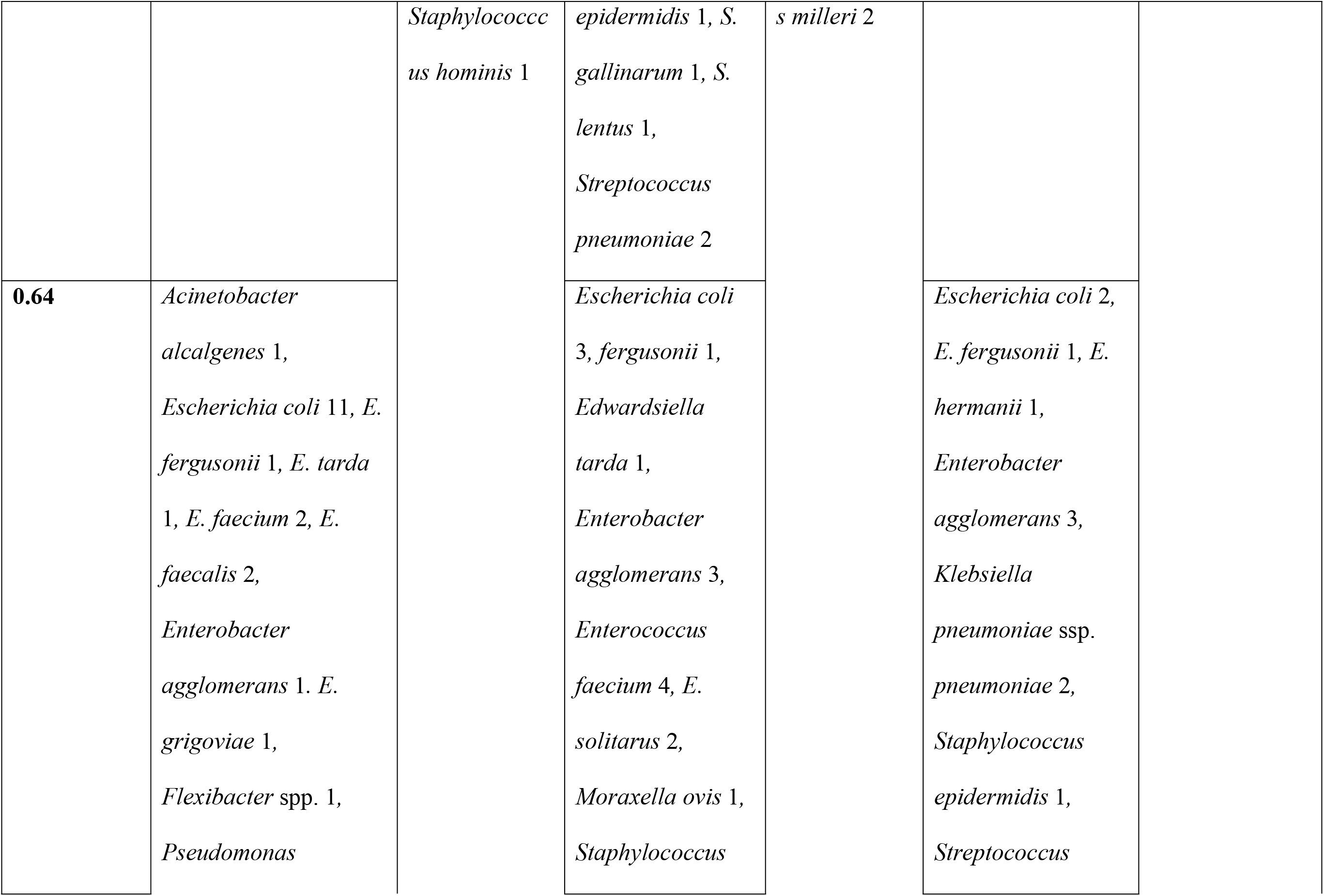
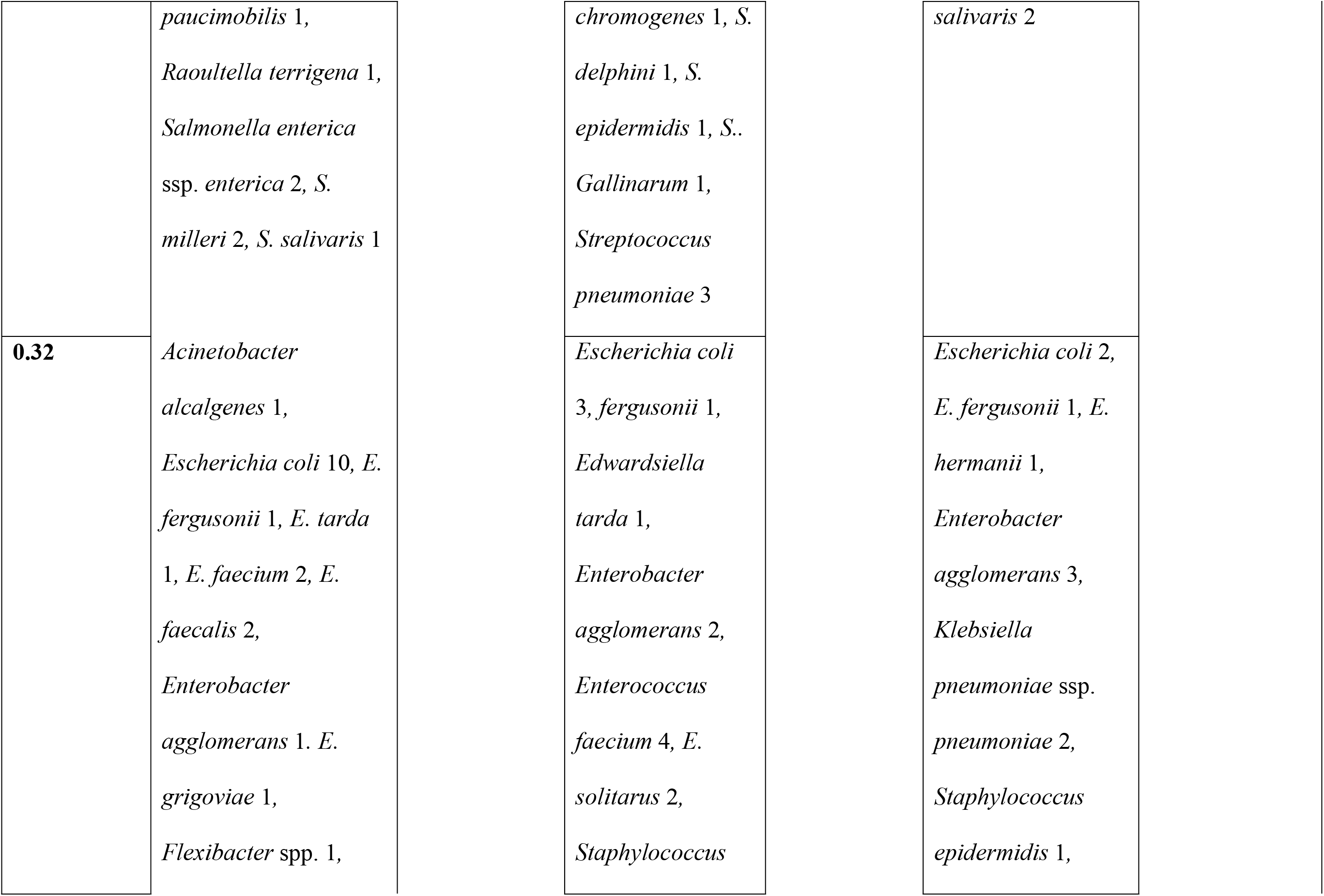
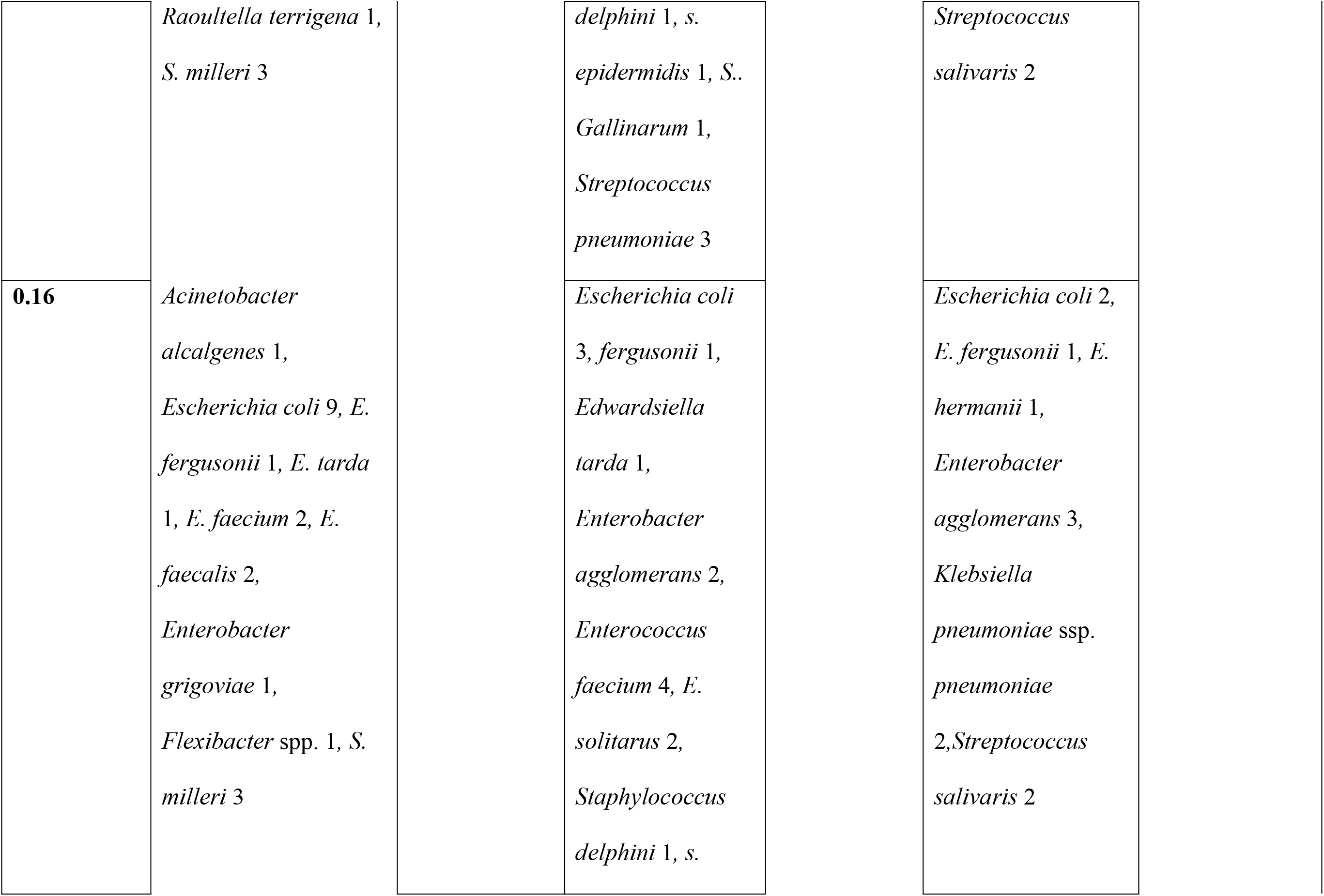
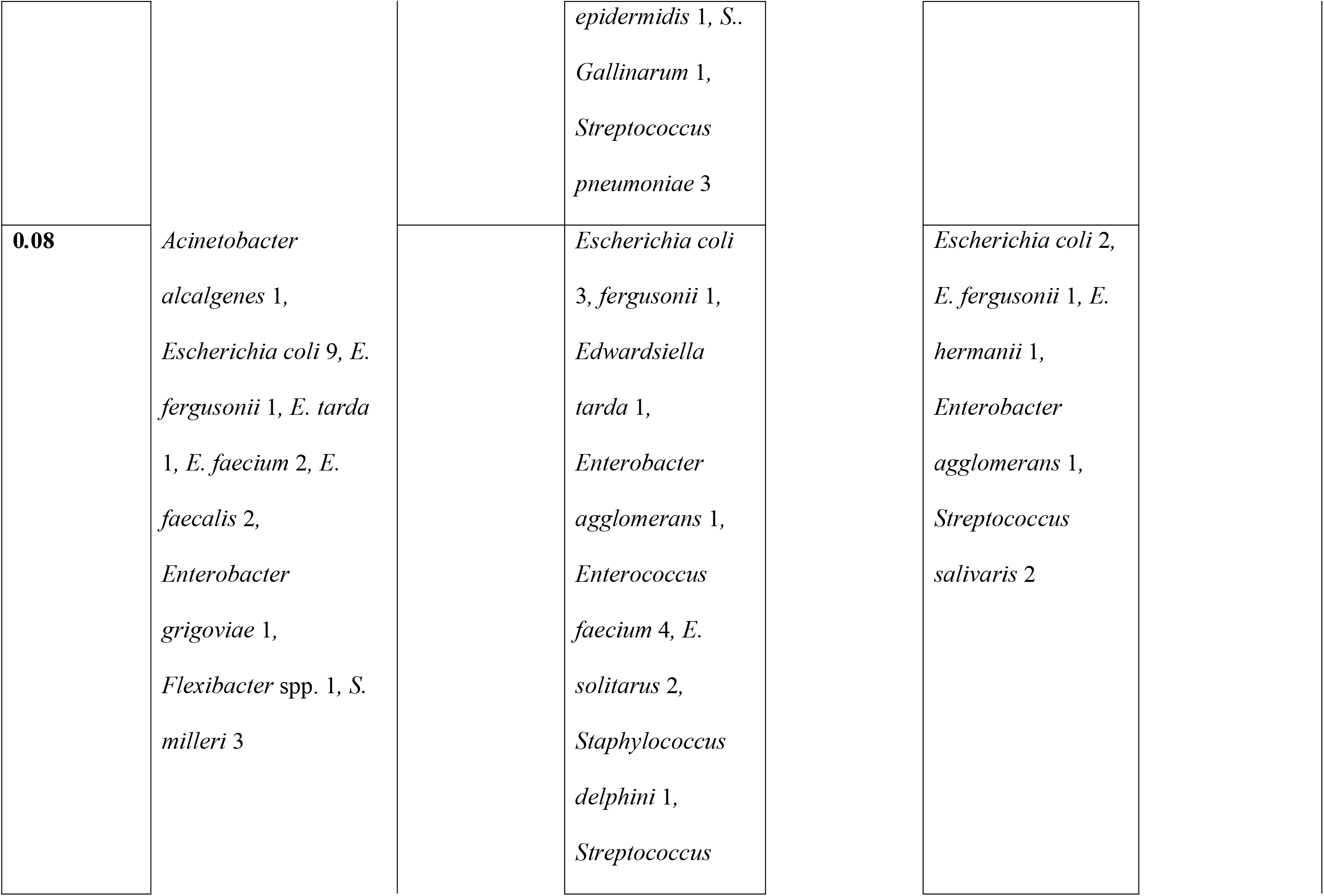
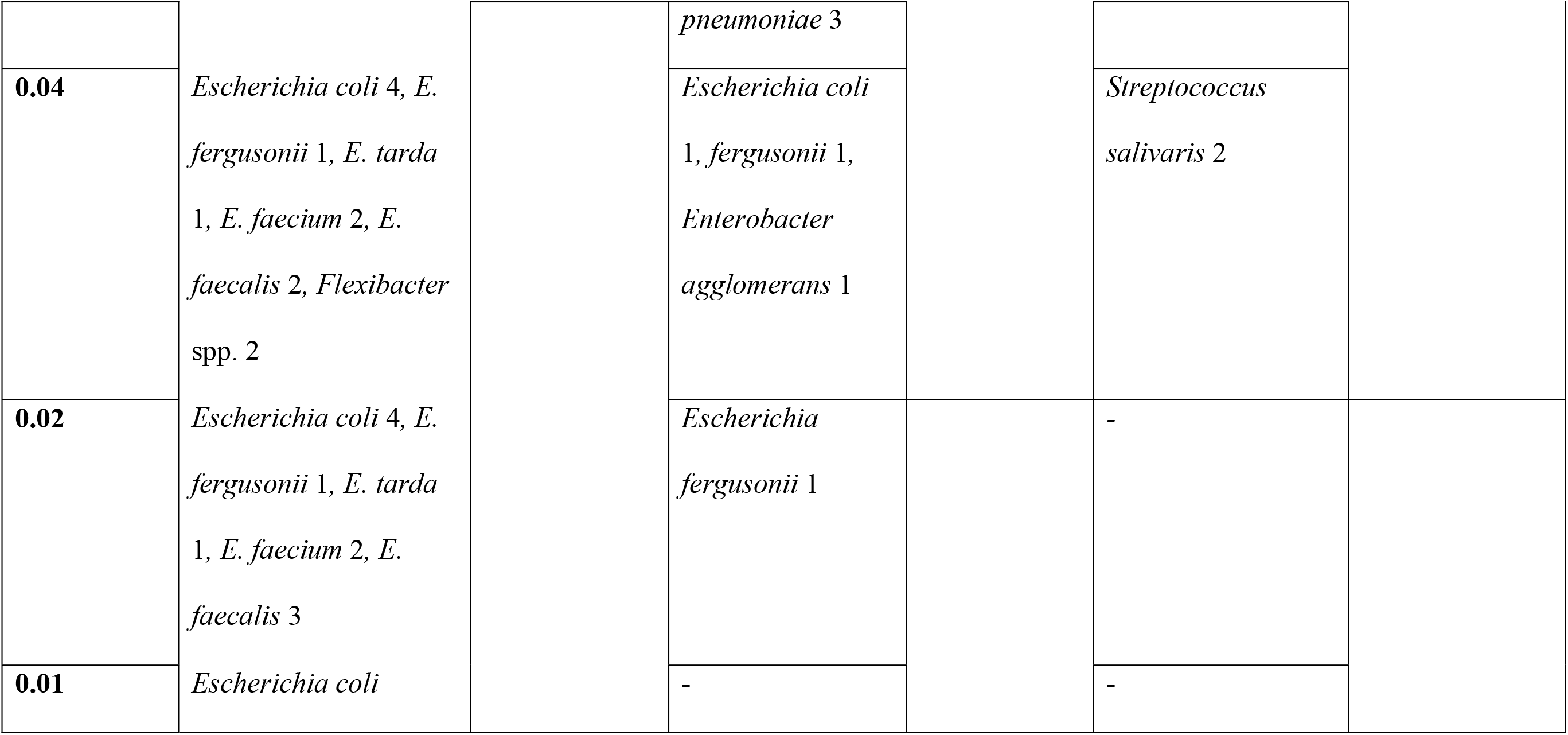
Synergistic effect of aspirin and paracetamol with minocycline and doxycycline antibiotic potential in terms of conversion of resistant bacteria to sensitive to the antibiotics.

## Results

A total of 293 strains of bacteria belonging to 32 genera (Tab. 1) were tested for MIC of aspirin, paracetamol, doxycycline and minocycline.

### MIC of aspirin

The MIC of aspirin was minimum (0.04 mg/ mL) for one strain each of *Staphylococcus xylosus* and *Streptococcus pyogenese* while it was maximum (10.24 mg/mL) for a few strains of *Burkholderia cepacia* (1/1), *Klebsiella pneumonia*e ssp. *pneumonia*e (1/12), *Pseudomonas aeruginosa* (3/6), *Staphylococcus aureus* (2/9), *S. capitis* ssp. *capitis* (1/3), *S. epidermidis* (6/10), *S. haemolyticus* (1/7) and *S. hominis* (2/5). Rest of strains tested had MIC of aspirin in between the two limits. The study indicated that about 1% solution of aspirin can stop the growth of bacteria.

### MIC of paracetamol

Except for one strain each of *Serratia grimaceae* (5.12 mg/mL) and *S, aureus* (5.12 mg/ mL) all the isolates grew in presence of 10.24 mg/ mL paracetamol, the maximum concentration used for testing.

### MIC of doxycycline

A *Pasteurella canis* strain was the most sensitive to doxycycline (MIC 0.125 μg/mL) while for another strain of *P. canis* MIC was 4 μg/mL. Of 293 strains tested for 46, 16, 25, 26, 24, 20, 20, 27, 12, 56, 7 and 13 strains had doxycycline MIC was 512 μg/mL, 256 μg/mL, 128 μg/mL, 64 μg/mL, 32 μg/mL, 16 μg/mL, 8 μg/mL, 4 μg/mL, 2 μg/mL, 1 μg/mL, 0.5 μg/mL and 0.25 μg/mL, respectively. A total of 116 (39.59%) strains were classified as sensitive (MIC ≤4 μg/mL) and 177 as resistant (MIC >4 μg/mL) to doxycycline.

### MIC of minocycline

Of 293 strains 8 strains had minocycline MIC equal to 0.125 μg/mL while 37, 8, 31, 14, 29, 23, 27, 25, 21, 24, 11, 34 and 1 strains had MIC equivalent to 0.25 μg/mL, 0.5 μg/mL, 1.0 μg/mL, 2.0 μg/mL, 4 μg/mL, 8 μg/mL, 16 μg/mL, 32 μg/mL, 64 μg/mL, 128 μg/mL, 256 μg/mL, 512 μg/mL and 1024 μg/mL, respectively. The most resistant strain was of *Hafnia alvei* with doxycycline MIC 512μg/mL. A total of 127 (43.34%) strain with MIC of minocycline ≤4 μg/mL were classified as sensitive and 166 as resistant (MIC >4 μg/mL) to minocycline.

The distribution of 293 strains of bacteria according to the MICs of doxycycline and minocycline was not normal and an erratic distribution curve was observed. The MICs of doxycycline and minocycline had a good positive correlation (r, 0.51; p 0.00001). There was no significant correlation was evident between MICs of doxycycline and aspirin but a strong correlation (r, 0.37; p 0.00001) was evident among MICs of aspirin and minocycline for different bacteria.

### Effect of aspirin on MIC of minocycline

A total of 164 strains of selected 24 genera (Tab. 2) were tested for determining MIC of minocycline in presence of 1.28 mg/mL, 0.64 mg/mL, 0.32 mg/mL, 0.16 mg/mL, 0.08 mg/mL, 0.04 mg/mL,, 0.02 mg/mL, and 0.01 mg/mL aspirin. Observations revealed that MIC of minocycline increased when tested in presence of aspirin for *Bacillus* species strains and was not affected for strains of *Burkholderia cepacia, Geobacillus stearothermophilus, Moelerella wisconsensis.* However, a significant reduction in MIC of minocycline was evident for strains of the rest of the 20 genera included in the study. In the study reduction in MIC was dependent on the concentration of aspirin in the media. The most affected strains having a reduction in minocycline MIC in presence of even 0.01 mg/ mL aspirin belonged to *Enterobacter* spp., *Escherichia coli, Hafnia alvei, Raoultella terrigena* and *Staphylococcus* species.

Of 164 strains 57 strains classified as minocycline-resistant (MIC >4 μg/mL) when tested for minocycline MIC in presence of 1.28 mg/mL, 0.64 mg/mL, 0.32 mg/mL, 0.16 mg/mL, 0.08 mg/mL, 0.04 mg/mL,, 0.02 mg/mL, and 0.01 mg/mL aspirin 32, 28, 23, 20, 20, 11, 10 and 1 (*E. coli*) strains become sensitive (MIC ≤4 μg/mL) to minocycline, respectively (Tab. 5). However, 25 minocycline-resistant strains retained their resistant to minocycline even in presence of aspirin. At near therapeutic plasma concentration (≤10 μg/mL) of aspirin only one minocycline-resistant *E. coli* strain turned sensitive to minocycline (MIC ≤4 μg/mL).

### Effect of aspirin on MIC of doxycycline

A total of 148 strains of selected 18 genera (Tab. 3) were tested for determining MIC of doxycycline in presence of 1.28 mg/mL, 0.64 mg/mL, 0.32 mg/mL, 0.16 mg/mL, 0.08 mg/mL, 0.04 mg/mL,, 0.02 mg/mL, and 0.01 mg/mL aspirin. Of the strains of 18 genera MIC of doxycyline got reduced for strains of 14 genera, increased for strains of *Aeromonas* spp. and *Bacillus* spp. and was not affected for strains of *R. terrigena* and *Paenibacillus* spp. The most affected strains to doxycycline in presence of aspirin even at 0.01 mg/ mL were of *Enterococcus* spp. and *Staphylococcus* spp.

Of the 148 strains 49 strains classified as doxycycline-resistant (MIC >4 μg/mL) on testing for doxycycline MIC in presence of 1.28 mg/mL, 0.64 mg/mL, 0.32 mg/mL, 0.16 mg/mL, 0.08 mg/mL, 0.04 mg/mL,, 0.02 mg/mL, and 0.01 mg/mL aspirin 30, 21, 18, 18, 15, 3 (*Escherichia coli, E. fergusonii*, *Enterobacter agglomerans*), 1 (*Escherichia fergusonii*) and 0 strains became sensitive (MIC ≤4 μg/mL) to doxycycline, respectively (Tab. 5). Of the 49 doxycycline-resistant strains 19 remained resistant to doxycycline even in presence of aspirin. At near therapeutic plasma concentration (≤10 μg/mL) of aspirin no doxycycline-resistant strains turned sensitive to doxycycline (MIC ≤4 μg/mL).

### Effect of paracetamol on MIC of doxycycline

A total of 112 strains of selected 11 genera (Tab. 4) were tested for determining MIC of doxycycline in presence of 1.28 mg/mL, 0.64 mg/mL, 0.32 mg/mL, 0.16 mg/mL, 0.08 mg/mL, 0.04 mg/mL,, 0.02 mg/mL, and 0.01 mg/mL acetoaminophen. The MIC of doxycycline reduced in presence of paracetamol for all but strains of *Moraxella* spp., *Proteus* spp. and *Pseudomonas* spp. The reduction in MIC of doxycycline was the most evident even at 0.01 mg/mL paracetamol for strains of doxycycline-sensitive (MIC ≤4 μg/mL) strains of *Erwnia* spp., *Escherichia* spp. and *Klebsiella pneumoniae* ssp. *pneumoniae*. However, the MIC of doxycycline for *Bacillus* ssp., *Erwinia* spp. and *Streptococcus* ssp. strains was not affected by presence of paracetamol at 1.28 mg/ mL, 0.64-1.28 mg/ mL and 1.28 mg/ mL, respectively.

Of the 112 strains 34 strains classified as doxycycline-resistant (MIC >4 μg/mL) on testing for doxycycline MIC in presence of 1.28 mg/mL, 0.64 mg/mL, 0.32 mg/mL, 0.16 mg/mL, 0.08 mg/mL, 0.04 mg/mL,, 0.02 mg/mL, and 0.01 mg/mL paracetamol 10, 11, 11, 10, 6, 1 (*Streptococcus salivaris*), 0 and 0 strains became sensitive (MIC ≤4 μg/mL) to doxycycline, respectively (Tab. 3). Of the 34 doxycycline-resistant strains 23 remained resistant even in presence of paracetamol. At near therapeutic plasma concentration (≤40 μg/mL) of paracetamol only one doxycycline-resistant strain of *S. salivaris* turned sensitive to doxycycline (MIC ≤4 μg/mL).

## Discussion

Though the MIC of doxycycline and minocycline for different bacteria to be classified as sensitive (16, 17) ranges between 0.25 μg/ mL (for *Neisseria gonorrhoeae*) to ≤4 μg/ mL (for members of Enterobacteriaceae and *Acinetobacter*), in the present study bacteria having MIC of doxycycline and minocycline ≤4 μg/ mL were considered sensitive for the therapeutic purposes. Because, infections with MIC ≤4 μg/ mL cab be treated effectively with doxycycline and minocycline as with therapeutic dosages it is an achievable plasma concentration of these drugs (16, 18). A total of 116 (39.59%) and 127 (43.34%) strains in the study were sensitive to doxycycline and minocycline, respectively. In the region of the study similar pattern of doxycycline and minocycline resistance is reported earlier too (19).

The erratic distribution of 293 strains of bacteria according to the MICs of doxycycline and minocycline might be attributed due to diversity among strains as they belonged to 32 genera with different sensitivity patterns to antibiotics. The strong positive correlation (r, 0.51; p 0.00001) among MICs of doxycycline and minocycline might due to the fact that both the antibiotics are of the same tetracycline class (15).

The least MIC of aspirin was detected 40 μg/ mL for one strain each of *Staphylococcus xylosus* and *Streptococcus pyogenese.* Plasma concentration of aspirin ranges between 4.9-8.9 μg/ mL in therapeutically applicable dosages and get converted to salicylate rapidly which may be in plasma with concentration of 42-62 μg/ mL and it may keep on increasing with chronic use of aspirin (20). Acute consumption of higher dosages more than 150 mg/kg may achieve higher plasma salicylate concentration but warrants an emergency detoxification treatment (21). The observations revealed that in therapeutically achievable concentration none of the bacterial strain in the study was treatable with aspirin.

At the maximum therapeutically achievable concentration (30 μg/ mL) of paracetamol in plasma, none of 293 strains tested in the study could be classified sensitive to paracetamol as the minimum MIC detected was 5.12 mg/ mL, that too only for one strain each of *Serratia grimaceae* and *S, aureus* i.e., for practical purposes paracetamol cannot be considered antimicrobial. In an earlier study, *S. aureus* has been shown to be sensitive to acetaminophen at 1.25 mg/ mL (22). The therapeutically useful plasma concentration of paracetamol may be between 10 to 20 μg/ mL and can be reached with oral as well intravenous use (23, 24). However, after a higher therapeutic dose on intravenous administration plasma concentration of paracetamol can be reached up to 30 μg/ mL. In supra-therapeutic toxic dosages plasma concentration of paracetamol is reported even up to 1500 μg/ mL (24).

In the present study only 42 (14.33%) strains were sensitive to aspirin at ≤1 mg/ mL concentration and none to paracetamol. Similarly, in an earlier study aspirin and paracetamol failed to contain growth of *Serratia, Bacillus* and *E. coli* strains at 1 mg/ mL concentration (25). Babik and coworkers (9) reported that aspirin failed to contain growth of *E. coli* and *S. epidermidis* at 1.5mg and 0.5 mg/ mL while paracetamol was not effective as antibacterial even at 3 mg/ mL concentration. In studies on *Campylobacter pylori* (now *Helicobacter pylori*) (26, 27) both aspirin and paracetamol are shown to be antibacterial in therapeutic dosages. In the present study neither strain of *Campylobacter* nor of *Helicobacter* were included thus the observation are not comparable. In a study (8), acetyl salicylic acid tested against *E. coli* and *S. aureus* and reported MIC equivalent to 2 mg/ mL.

In the present study MIC of doxycycline increased in presence of aspirin for strains of *Aeromonas* spp. and *Bacillus* spp. and MIC of minocycline also increased in presence of aspirin for *Bacillus* spp. strains. Similar observations are reported by Hadera and coworkers (8) while evaluating aspirin in combination with ciprofloxacin and benzylpenicillin against *E. coli* and *S. aureus* and reported antagonistic drug interaction. However, in the present study, none of the *E. coli* and *S. aureus* and strains of other genera showed antagonism between aspirin / paracetamol and minocycline/ doxycycline.

In the study, aspirin converted a minocycline-resistant *E. coli* and doxycycline-resistant *S. salivaris* strains to sensitivity even at 10μg/ mL levels of aspirin. However, a total of 32 out of 57 minocycline-resistant strains became sensitive to minocycline and 30 of 42 doxycycline-resistant strain reverted to be sensitive to doxycycline in presence of 1280μg/ mL levels of aspirin. Earlier studies (28) reported that aspirin, sodium salicylate and sodium benzoate reverted colistin-resistant Enterobacteriaceae and *P. aeruginosa* to sensitivity. Chan and coworkers also (29) reported synergy between aspirin and antibiotics such as cefuroxime and chloramphenicol when administered against methicillin-resistant *Staphylococcus aureus*.

Observations indicated that synergy between aspirin and doxycycline/ minocycline might be strain-dependent. Observations are in concurrence to earlier studies. Ahmed and coworkers (7) reported no-synergistic interaction between aspirin and ß-lactam antibiotics (ampicillin, amoxicillin, amoxicillin + clavulanic acid, cephalexin and cefotaxime) against *P. aeruginosa* and *K. pneumoniae* strains but reported synergistic action of aspirin with amoxicillin, cefotaxime, augmentin, gentamicin and ciprofloxacin against *E. coli* strains (6).

Paracetamol also showed synergistic activity with doxycycline and 11 of the 34 doxycycline-resistant strains turned sensitive when tested in presence of paracetamol ≤1.28 mg/ mL. Though there are several studies on interactions of NSAIDs with antibiotics, the interaction of the most commonly prescribed antipyretic paracetamol are rarely reported (30).

The synergy or adjuvant activity of aspirin and paracetamol with doxycycline and minocycline was strain-specific, within the same species of bacteria both revertible and non-revertible strains were present (Tab. 3). Thus, it is apparent that synergy between NSAIDs and antibiotics is modulated by bacterial factors rather than just combination of the two drugs. Further targeted studies may reveal genes/ factors responsible for synergy or no-synergy or antagonism between NSAIDs and antibiotics to help understanding the phenomenon and finally in development of a strategy to mitigate AMR and convert partly inefficient antibiotics to therapeutically useful ones, that is revival of outdated antibiotics.

The study concluded that neither aspirin nor paracetamol have potential antibacterial activity at therapeutic dose levels. However, aspirin at 10.24 mg/ mL (1.024%) concentration inhibited all 293 strains in the study while paracetamol could not inhibit any but one strain each of *S. aureus* and *Serratia grimaceae* at the same concentration. Though aspirin cannot be used at the bacteria inhibiting concentration (1.024%) internally, its topical use either as lotion or ointment may be promising as a broad spectrum antibacterial even against the strains showing resistance to antibiotics. Further, the presence of aspirin converted 32 of 57 minocycline-resistant strains and 30 of 42 doxycycline-resistant strains to sensitivity. The synergy between aspirin even at 0.1% concentration level with minocycline and doxycycline can be utilized for the formulation of topical antibacterials. In contrast, paracetamol appeared almost ineffective as antibacterial even at >1% concentration and was also not as efficient adjuvant as aspirin to minocycline and doxycycline to increase their antibacterial activity. The study also indicated the need for more studies to reveal the mechanism underlying the interaction and synergy between NSAIDs and antibiotics.

## Acknowledgements

The author is thankful to Mr. HC Joshi, Mr. G Tiwari, Mr. Pratap Singh, Mr. Laiqur Rahman and Mr. Ashok Kumar, Division of Epidemiology, ICAR-Indian Veterinary Research Institute, Izatnagar for their consistent technical assistance. The research work was supported by grants received from CAAST-ACLH (NAHEP/CAAST/2018-19).

## References

1. Gajdács M. 2020. Non-antibiotic pharmaceutical agents as antibiotic adjuvants. Acta Biologica Szegediensis 64:17–23.

2. Miró-Canturri A, Ayerbe-Algaba R, Smani Y. 2019. Drug repurposing for the treatment of bacterial and fungal infections. Front Microbiol 10:41 doi: 10.3389/fmicb.2019.00041.

3. Khan S, Dhama K, Patel SK, Pathak M, Tiwari R, Singh BR, Shah R, Bonilla-Ardana K, Rodriguez-Morales AJ, Leblebicioglu H. 2020. Ivermectin, a new candidate therapeutic against SARS-CoV-2/COVID-19. Annals Clin Microbiol Antimicrobials 19: 23 doi: 10.1186/s12941-020-00368-w.

4. Lagadinou M, Onisor MO, Rigas A, Musetescu D, Gkentzi D, Assimakopoulos SF, Panosand G. Markos Marangos M. 2020. Antimicrobial properties on non-antibiotic drugs in the era of increased bacterial resistance. Antibiotics 9:107 doi:10.3390/antibiotics9030107.

5. Leão RP; Cruz JV, da Costa GV, Cruz JN, Ferreira EFB, Silva RC, de Lima LR, Borges RS, dos Santos GB, Santos CBR. 2020. Identification of new rofecoxib-based cyclooxygenase-2 inhibitors: a bioinformatics approach. Pharmaceuticals 13:209.

6. Ahmed EFR, Abd El-Baky RM, Ahmed ABF, Fawzy NG, Aziz NA, Gad GFM. 2016. Evaluation of antibacterial activity of some non-steroidal anti-inflammatory drugs against *Escherichia coli* causing urinary tract infection. African J Microbiol Res 10:1408–1416.

7. Ahmed EFR, El-baky MA, Ahmed ABF, Waly NG, Gad GFM. 2017. Antibacterial activity of some non-steroidal anti-inflammatory drugs against bacteria causing urinary tract infection. Am J Infect Dis Microbiol 5:66–73.

8. Hadera M, Selam Mehari S, Saleem Basha S, Nebyu D, Amha ND, Berhane Y. 2018. Study on antimicrobial potential of selected non-antibiotics and its interaction with conventional antibiotics. UK. J. Pharm. Biosci. 6:1–7

9. Babik M. 2021. Utilizing the antimicrobial effects of antipyretics through synergistic combination with other FDA approved drugs. Research Gate doi: 10.13140/RG.2.2.10337.63841.

10. Chow JH, Khanna AK, Kethireddy S, Yamane D, Levine A, Jackson A, McCurdy MT. et al. 2021. Aspirin use is associated with decreased mechanical ventilation, intensive care unit admission, and in-hospital mortality in hospitalized patients with Coronavirus Disease 2019. Anesthesia Analgesia 132:930–941. doi: 10.1213/ANE.0000000000005292.

11. Ministry of Health and Family Welfare, Govt. of India. 2021. Revised guidelines for home isolation of mild /asymptomatic COVID-19 cases. https://www.mohfw.gov.in/pdf/RevisedguidelinesforHomeIsolationofmildasymptomaticCOVID19cases.pdf

12. Yates PA, Newman SA, Oshry LJ, Glassman RH, Leone AM, Reichel E. 2020. Doxycycline treatment of high-risk COVID-19-positive patients with comorbid pulmonary disease. Ther Adv Respir Dis 14:1753466620951053 doi: 10.1177/1753466620951053.

13. Francini E, Miano ST, Fiaschi AI, Francini G. 2020. Doxycycline or minocycline may be a viable treatment option against SARS-CoV-2. Med Hypotheses 144:110054. doi:10.1016/j.mehy.2020.110054

14. Singh H, Kakkar AK, Chauhan P. 2020. Repurposing minocycline for COVID-19 management: mechanisms, opportunities, and challenges. Expert Rev Anti Infect Ther 18:997–1003 doi: 10.1080/14787210.2020.1782190.

15. Singh BR. 2009. Labtop for microbiology laboratory. Lamber t Academic Publishing, AG & Co. KG, Berlin, Germany.

16. CLSI. 2014. Performance standards for antimicrobial disk susceptibility tests, in: Institute. Clinical and Laboratory Standards Institute, Wayne, Pennsylvania.

17. CLSI, 2015. Methods for Antimicrobial Dilution and Disk Susceptibility Testing of Infrequently Isolated or Fastidious Bacteria. Clinical and Laboratory Standards Institute, Wayne, USA.

18. Cunha BA, Domenico P, Cunha C. 2000. Pharmacodynamics of doxycycline. Clin Microbiol Infect 6: 270–273. https://doi.org/10.1046/j.1469-0691.2000.00058-2.x

19. Singh BR. 2018. ESKAPE Pathogens in animals and their antimicrobial drug resistance pattern. J Dairy Vet Anim Res 7(3):10 doi: 10.19080/JDVS.2018.07.555715.

20. National Center for Biotechnology Information. 2021. PubChem compound summary for CID 2244, aspirin. https://pubchem.ncbi.nlm.nih.gov/compound/Aspirin.

21. Chyka PA, Erdman AR, Christianson G, Wax PM, Booze LL, Manoguerra AS, Caravati EM, Nelson LS, Olson KR, Cobaugh DJ, Scharman EJ, Woolf AD, Troutman WG. 2007. Salicylate poisoning: an evidence-based consensus guideline for out-of-hospital management. Clin Toxicol (Phila) 45:95–131 doi: 10.1080/15563650600907140.

22. Al-Janabi AA. 2010. *In vitro* antibacterial activity of ibuprofen and acetaminophen. J Glob Infect Dis 2:105–108. doi:10.4103/0974-777X.62880

23. Rumack BH. 1978. Aspirin versus acetaminophen: a comparative view. Pediatrics 62(5 Pt 2 Suppl):943–946.

24. Medsafe. 2012. New Zealand Data Sheet. Paracetamol AFT inf. https://www.medsafe.govt.nz/profs/Datasheet/p/paracetamolAFTinf.pdf

25. Lawal A, Obaleye JA. 2008. Synthesis, Characterization and antibacterial activity of aspirin and paracetamol-metal complexes. Nigerian Soc Exp Biol https://tspace.library.utoronto.ca/handle/1807/44715

26. Caselli M, Pazzi P, LaCorte R, Aleotti A, Trevisani L, Stabellini G. 1989. *Campylobacter*-like organisms, nonsteroidal anti-inflammatory drugs and gastric lesions in patients with rheumatoid arthritis. Digestion 44:101–104.

27. Wang WH, Wong WM, Dailidiene D, Berg DE, Gu Q, Lai KC, Lam SK, Wong BC. 2003. Aspirin inhibits the growth of *Helicobacter pylori* and enhances its susceptibility to antimicrobial agents. Gut 52:490–495.

28. Malla CF, Mireles NA, Ramírez AS, Poveda JB, Tavío MM. 2020. Aspirin, sodium benzoate and sodium salicylate reverse resistance to colistin in Enterobacteriaceae and *Pseudomonas aeruginosa*. J Antimicrob Chemother 75:3568–3575. doi: 10.1093/jac/dkaa371

29. Chan EWL, Yee YZ, Raja I, Yap JKY. 2017. Synergistic effect of non-steroidal anti-inflammatory drugs (NSAIDs) on antibacterial activity of cefuroxime and chloramphenicol against methicillin-resistant *Staphylococcus aureus*. J Glob Antimicrob Resist 10:70–74.

30. InformedHealth.org. 2016. Cologne, Germany: Institute for Quality and Efficiency in Health Care (IQWiG); 2006-. Using medication: The safe use of over-the-counter painkillers. https://www.ncbi.nlm.nih.gov/books/NBK361006/

